# Differential cell type-specific function of the aryl hydrocarbon receptor and its repressor in diet-induced obesity and fibrosis

**DOI:** 10.1101/2024.02.22.581524

**Authors:** Frederike J. Graelmann, Fabian Gondorf, Yasmin Majlesain, Birte Niemann, Katarina Klepac, Marlene Gottschalk, Michelle Mayer, Irina Iriady, Philip Hatzfeld, Sophie K. Lindenberg, Klaus Wunderling, Christoph Thiele, Zeinab Abdullah, Wei He, Karsten Hiller, Kristian Händler, Marc D. Beyer, Thomas Ulas, Alexander Pfeifer, Charlotte Esser, Heike Weighardt, Irmgard Förster, Laia Reverte-Salisa

## Abstract

The aryl hydrocarbon receptor (AhR) is a ligand-activated transcription factor regulating xenobiotic responses as well as physiological metabolism. Dietary AhR ligands activate the AhR signaling axis in the intestine and throughout the organism, whereas AhR activation is negatively regulated by the AhR repressor (AhRR). While AhR-deficient mice are known to be resistant to diet-induced obesity (DIO), we here demonstrate that AhRR deficiency also leads to a robust, but not as profound protection from DIO and hepatosteatosis. Under conditions of DIO, AhRR^-/-^ mice did not accumulate TCA cycle intermediates in the circulation in contrast to wild-type (WT) mice, indicating protection from metabolic dysfunction. This effect could be mimicked by dietary supplementation of AhR ligands in WT mice. Because of the predominant expression of the AhRR in myeloid cells, AhRR-deficient macrophages were analyzed for changes in metabolism and showed major metabolic alterations regarding oxidative phosphorylation and mitochondrial activity as well as increased expression of genes involved in *de novo* lipogenesis and mitochondrial biogenesis. Mice with a genetic deficiency of the AhRR in myeloid cells did not show alterations in weight gain after high fat diet (HFD) but demonstrated ameliorated liver damage compared to control mice. Further, deficiency of the AhR in myeloid cells also did not affect weight gain but led to enhanced liver damage and adipose tissue fibrosis compared to controls. Although conditional ablation of either the AhR or AhRR in myeloid cells did not recapitulate the phenotype of the global knockout, our findings suggest that enhanced AhR signaling in myeloid cells deficient for AhRR protects from diet-induced liver damage and fibrosis, whereas myeloid cell-specific AhR deficiency is detrimental.

## 1. Introduction

The aryl hydrocarbon receptor (AhR) is a sensor of polyaromatic chemicals, including environmental toxins (xenobiotic ligands), phytochemicals, and microbial or endogenous metabolites (natural ligands) [1–3]. Upon ligand binding and activation, AhR translocates from the cytoplasm into the nucleus for regulation of target gene expression. Transcriptional regulation of gene expression through the AhR has a major impact on immunity [4–6] as well as cellular and systemic metabolism [7–9]. While high-dose exposure with 2,3,7,8-tetrachlorodibenzo-p-dioxin (TCDD) causes hepatic steatosis [10–12] and wasting syndrome [13,14], chronic low dose exposure to industrial pollutants has been implicated in the development of obesity and insulin resistance [15–18]. Mice with a complete deficiency of the AhR [19] or those bearing a low-affinity AhR mutant [8] have been shown to be protected from hepatic steatosis and DIO. Such resistance was not or only partially seen in animals with a hepatocyte-specific AhR deficiency [20], while mice with an adipocyte-specific knockout of the AhR show a sexually dimorphic effect on weight gain when fed a HFD [15,21]. These findings indicate an important yet cell type-specific function of the AhR that may either enhance or protect from DIO and metabolic dysregulation. Expression of a constitutively active mutant of the AhR under control of the fatty acid binding protein (FABP) promoter in the liver caused hepatic steatosis but protected the animals from DIO due to enhanced expression of CD36 and production of the hepatokine fibroblast growth factor 21 (FGF21) as a direct transcriptional target of AhR [12,22]. Furthermore, several key metabolic enzymes involved in lipid metabolism were found to be transcriptionally regulated through the AhR [23,24].

Considering the major impact of AhR activation on cellular metabolism as well as immune regulation by xenobiotic and endobiotic ligands [1,3,25,26], the AhR signaling pathway is tightly regulated through feedback inhibition. Enzymes of the Cytochrome p450 monooxygenase (CYP) family, such as CYP1A1, CYP1A2, and CYP1B1, are strongly induced by AhR activation and rapidly metabolize AhR ligands leading to a shut-down of this signaling pathway [4,27,28]. Two other target genes of the AhR encode the TCDD-inducible poly (ADP-ribose) polymerase (TiPARP) and the AhRR, which turn down AhR signaling by different means, either through enhanced ribosylation-mediated protein degradation of the AhR or by competition for binding to AhR nuclear translocator (ARNT), respectively [1,29–31].

We previously generated an *Ahrr* reporter allele in mice, in which the enhanced green fluorescent protein (EGFP) is expressed under the control of the endogenous *Ahrr* promoter. Using these AhRR/EGFP reporter mice, we demonstrated that the AhRR is mainly expressed in immune cells of the skin and intestinal mucosa, including myeloid cells, innate lymphoid cells, and several T cell subsets [32]. No notable expression of the AhRR could be detected in intestinal epithelial cells at steady state or following systemic AhR stimulation, whereas other AhR target genes are known to be expressed at this site [4,33]. Thus, the AhRR appears to play a predominant role in regulating AhR activation in a cell type- or organ-specific manner, in particular in hematopoietic cells [32,34,35]. Functionally, AhRR-dependent regulation of AhR activity was shown to be important for the balancing of cytokine production by myeloid cells and T cells in the context of inflammation [32]. Recently, it was shown that AhRR deficiency decreases intestinal intraepithelial lymphocyte (IEL) numbers in an oxidative stress-dependent manner, facilitating intestinal infection and inflammation [36].

Tissue-resident immune cells have been demonstrated to be essential regulators of lipid metabolism, energy consumption, and thermogenesis in adipose tissues [37–42]. Therefore, the regulation of AhR function through the AhRR, which is predominantly expressed in immune cells, may represent an important rheostat of lipid metabolism. Macrophages, in particular, are known to control homeostasis in metabolic organs such as the liver [43–45] and adipose tissue [46–48], depending on their activation status [41,42,49].

In the present study, we evaluated the relevance of the AhRR as well as the dietary induction of AhR signaling for systemic and cellular metabolism. We also determined the sensitivity of complete and myeloid cell-specific AhRR-deficient mice to DIO in direct comparison to AhR deficiency.

## 2. Results

### 2.1 AhRR-deficient mice are partially resistant to diet-induced obesity

Here we asked whether enhanced AhR activation by deletion of the AhRR, a negative regulator of AhR signaling, might impact the induction of DIO and the resulting disturbances in metabolism. Activation of the AhR by xenobiotic or endogenous ligands is likely to contribute to DIO, as AhR-deficient mice or mice expressing the low-affinity AhR are protected from diet-induced weight gain and liver steatosis [8,19]. In line, the inhibition of AhR activation by CH-223191 led to the same effect [50]. When global AhRR-deficient mice were fed with HFD for 14 weeks, their weight gain was significantly lower than that of WT control mice (Fig. 1A). Blood glucose levels were significantly lower in AhRR-deficient mice compared to WT mice after glucose and insulin stimulation (Fig. 1B). In addition, total serum cholesterol levels were significantly reduced in mice lacking the AhRR (Fig. 1C). In line with these results, AhRR^-/-^ HFD-fed mice had higher O_2_ consumption at room temperature (Fig. 1D). As expected, the total fat mass was increased in HFD WT compared to CD mice, but this weight gain was strongly decreased in HFD AhRR^-/-^ compared to WT HFD mice (Fig. 1E). In alignment with the total fat mass, the weight of gonadal white adipose tissue (WATg) was increased by 198% after HFD feeding, while AhRR-deficient mice had 53.4% less WATg compared to WT HFD mice (Fig. 1F). Moreover, this reduction in WATg weight was accompanied by a reduced adipocyte size in mice lacking AhRR (Fig. 1G). This indicates that deficiency of the AhRR protects from weight gain and enlargement of adipose tissue after HFD feeding in mice. Another hallmark of DIO is liver steatosis. Here, liver weight was increased in WT mice after HFD feeding. In contrast, the liver weight was unaltered in HFD-fed AhRR-deficient mice compared to CD-fed mice (Fig. 1H). Histological analysis revealed that lipid droplet accumulation in the liver was reduced in HFD-fed AhRR-deficient mice. Also, a reduced level of neutral lipids in the liver tissue was demonstrated by reduced staining intensity of Oil Red O (ORO) (Fig. 1I-J). Accordingly, while serum levels of alanine aminotransferase (ALT) were enhanced in HFD-fed WT mice, indicating liver damage, this was not observed in HFD-fed AhRR-deficient mice (Fig. 1K).

**Figure 1:**
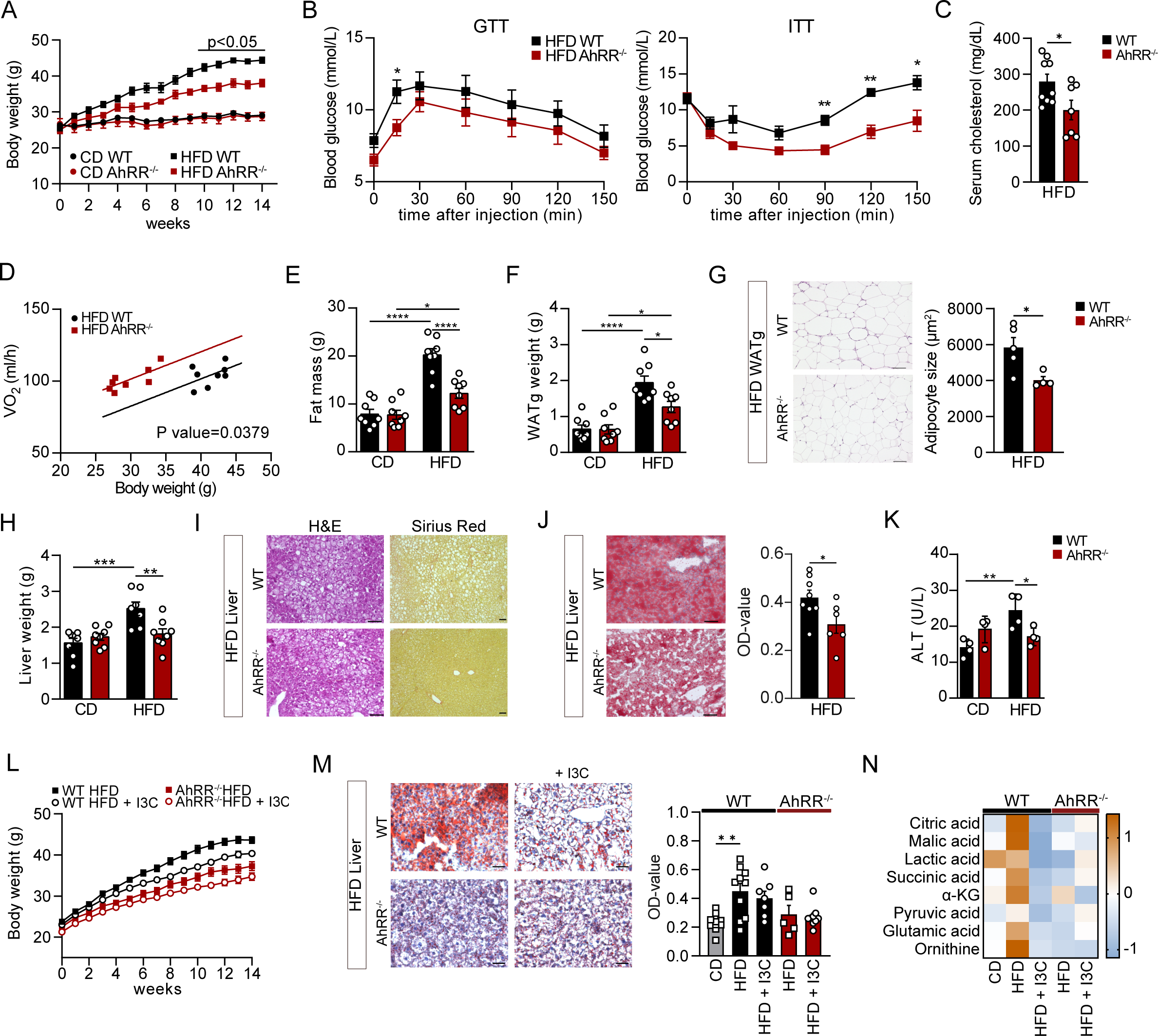
AhRR-deficient mice are partially resistant to diet-induced obesity. (**A**) Body weight of wild-type (WT) and AhRR-deficient mice fed CD or HFD for 14 weeks; n= 6-8 (WT), n=5 (AhRR^-/-^). (**B**) Blood glucose levels during a glucose tolerance test -GTT- (left panel) and insulin tolerance test -ITT- (right panel) after 12 weeks on HFD; n=10-11 (GTT), n=5 (ITT). (**C**) Total serum cholesterol levels after 14 weeks on HFD; n=7-9. (**D**) Analysis of covariance (ANCOVA) (non-linear fit) of oxygen consumption/body weight (BW); n=8. (**E**) Total body fat mass assessed by NMR; n=8. (**F**) WATg weight; n=8. (**G**) Representative H&E staining. Scale bars, 80 µm (left panel). Right, quantification of adipocyte size of the WATg of HFD mice; n=4-5. (**H**) Liver weight; n=7-8. (**I**) Representative H&E (left panel) and Sirius Red (right panel) staining of liver sections. Scale bars, 100 µm. (**J**) Representative Oil Red O (ORO) and Hemalaun staining of liver sections of WT and AhRR-deficient mice after HFD. Scale bars, 80 µm (left panel). Right, quantification of ORO stain, n=6-8. (**K**) Serum Alanine transaminase (ALT) concentration of WT and AhRR-deficient mice fed either CD or HFD for 14 weeks; n=4. (**L**) Body weight of WT and AhRR-deficient mice fed a HFD supplemented with or without indole-3 carbinole (I3C) for 14 weeks; n=15-20 (WT) and n=14-15 (AhRR^-/-^). (**M**) Representative ORO and Hemalaun staining of liver sections of WT and AhRR^-/-^ mice after HFD ± I3C. Scale bars, 100 µm (left panel). Right, ORO staining quantification of liver sections; n=7-10 (WT), n= 5-9 (AhRR^-/-^). (**N**) z-scored intensity values of representative TCA-metabolites found in plasma of mice after 14 weeks of CD or HFD ± I3C; n=3-5. *p < 0.05, **p ≤0.01, ***p ≤ 0.001. Significance was determined using unpaired two-tailed t-tests (A, B, C, G and J) and one-way analysis of variance (ANOVA) with Tukey’s multiple-comparison test (E, F, H, K and M). Data are mean ± s.e.m.

As activation of the AhR by feeding indole-3 carbinol (I3C) protects mice from DIO affecting body weight gain and lipid accumulation [51], we wondered whether additional AhR activation with I3C affects DIO in AhRR-deficient mice. As expected, WT mice fed HFD supplemented with 2 g/kg I3C gained less weight compared to WT mice fed HFD only (Fig. 1L). AhRR-deficient mice again had reduced weight gain, with no notable further reduction of weight gain after feeding the I3C-supplemented HFD (Fig. 1L). The lower weight gain in WT mice fed I3C-supplemented HFD was only marginally reflected by a reduced accumulation of lipid droplets in the liver. AhRR-deficient mice, in contrast, already show reduced lipid buildup after feeding HFD without further reduction after I3C supplementation (Fig. 1M). Similar to published data [52], metabolomic analysis of the plasma revealed an increase in TCA cycle intermediates in WT HFD mice hinting to metabolic dysfunction upon 14 weeks of HFD, which was absent in HFD AhRR-deficient mice and in both groups fed an I3C-supplemented HFD (Fig. 1N). As expected, AhR-deficient mice only moderately gained weight after HFD feeding, while supplementation with I3C did not influence body- or liver weight gain in these mice, indicating that the observed effect of I3C supplementation is indeed AhR-dependent (Fig. S1A). In line, I3C supplementation did not elicit any changes in the plasma metabolome of AhR^-/-^ mice (Fig. S1B). In conclusion, the above findings indicate that metabolic alterations induced by DIO can be reverted by activation of the AhR.

### 2.2 AhRR-deficient macrophages display an altered metabolic and gene expression profile

Since AhRR expression is most prominent in immune cells, such as myeloid cells [32,53], we assessed whether AhRR expression modulates the metabolism and gene expression profiles in macrophages. For this purpose, we compared the transcriptome profiles using WT and AhRR-deficient long-term Fetal Liver Macrophage (FLiM) cell lines [54]. AhRR^-/-^ FLiM cells showed a profound alteration in their transcriptional profile in comparison to WT FLiM cells (Fig. 2A-B, S2A) with a total of 672 differentially expressed genes (DEGs) (312 up, 360 down). Hallmark Gene Set enrichment analysis (GSEA) was performed on result of the DEG analysis comparing AhRR^-/-^ vs WT FLiM cells. This analysis revealed an enrichment for Myc- and mTORC1 pathways known to control protein synthesis, metabolism, and mitochondrial biogenesis [55,56] in AhRR^-/-^ compared to WT FLiM cells (Fig. 2C). Markedly, genes involved in fatty acid and lipid mediator synthesis (*Acaca, Elovl6, Acly, Ch25h*) were also higher expressed in AhRR^-/-^ vs WT FLiM cells (Fig. 2B, S2A, Supplementary Table 1 (GSEA)). Pathways influencing the TNF and interferon response were negatively enriched in AhRR^-/-^ FLiM cells, alongside genes associated with adipogenesis and lipoprotein metabolism (*Apoc1, Apoc4*) and lipid storage (*Plin2, Abca1*) (Fig. 2B, S2A). These findings highlight the involvement of the AhR/AhRR signaling axis in metabolic processes on a cellular level, regulating key cellular programs such as protein synthesis and immunometabolism.

**Figure 2:**
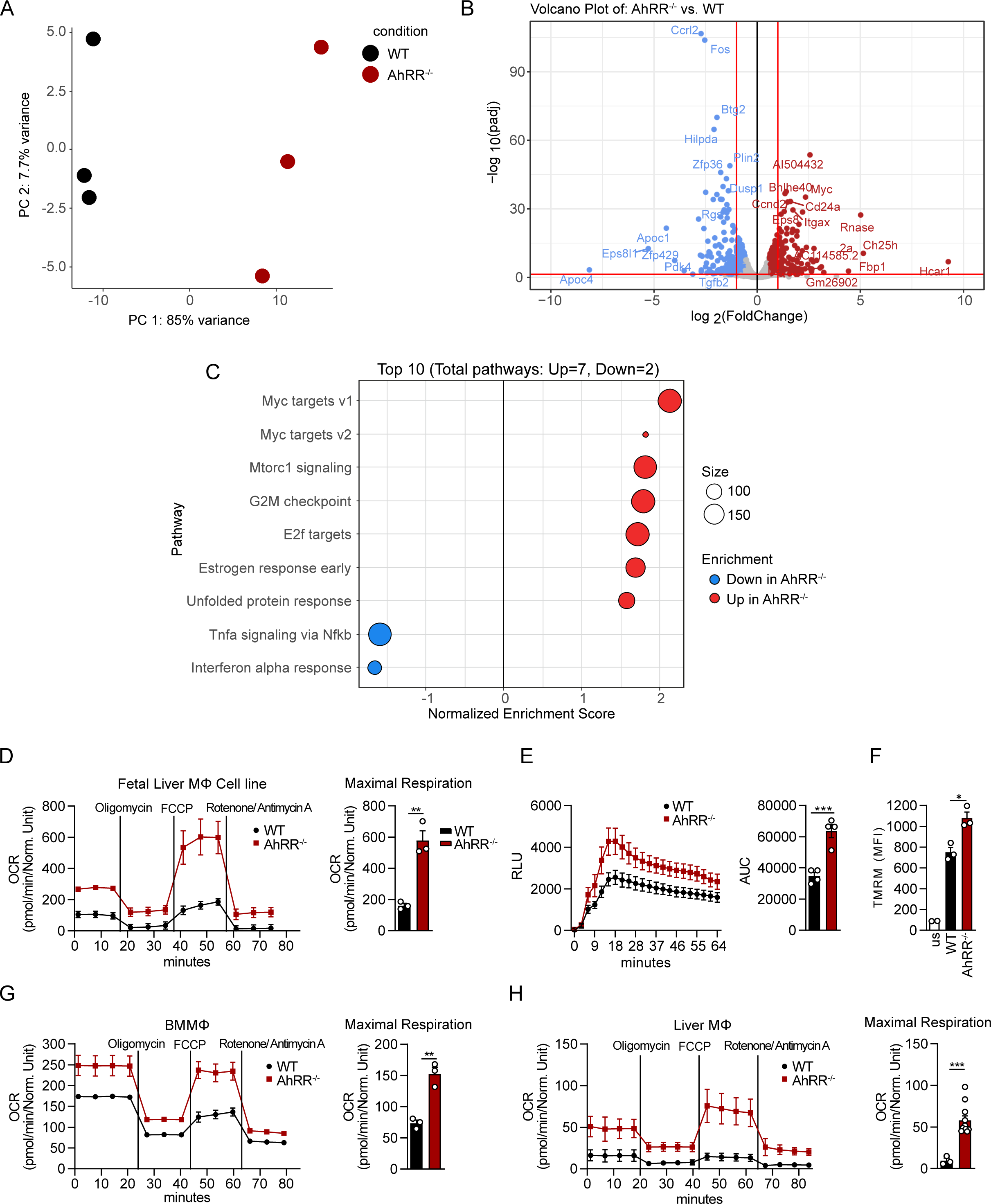
AhRR-deficient macrophages display an altered metabolic and gene expression profile. (**A**) Principal Component Analysis (PCA) plot depicting relationship of all samples based on dynamic gene expression of all genes comparing WT and AhRR^-/-^ Fetal Liver MΦ (FLiM) cells (n=3) and (**B**) Volcano plot depicting fold changes (FC) and FDR-adjusted p values comparing WT and AhRR^-/-^ FLiM cells. Differentially expressed up- (red) and downregulated genes (blue) are shown and selected genes are highlighted. (**C**) GSEA results based on the ranked gene list between AhRR^-/-^ vs WT FLiM cells, n=3. Only significant enriched terms are visualized by the Normalized enrichment scores (NES) and the enriched terms based on the Hallmark DB, n=3. (**D**) Mitochondrial stress test analysis of FLiM cells of WT and AhRR^-/-^ mice using a Seahorse XFe96 analyzer. Oxygen consumption rate (OCR) over time (left panel) and quantification of the maximal OCR (right panel); n=3. (**E**) ROS production of FLiM cells after stimulation with zymosan over time (left panel) and quantification depicted as area under curve (AUC, right panel); n=4. (**F**) Flow cytometric measurement of mitochondrial membrane potential using Tetramethylrhodamine Methyl Ester (TMRM), depicted as MFI; n=3. us = unstained control, n=2. (**G**) Mitochondrial stress test analysis in bone marrow-derived macrophages (BMMΦ) and (**H**) primary liver macrophages of WT and AhRR^-/-^ mice. OCR over time (left panel) and quantification of the maximal OCR (right panel); For (G), n= 3 and for (H), n=3 (WT), n=10 (AhRR^-/-^). Seahorse and ROS data show mean ± s.e.m. of three technical replicates from one representative experiment. *p < 0.05, **p ≤0.01, ***p ≤ 0.001. Significance was determined using unpaired two-tailed t-tests. Data are mean ± s.e.m.

To directly assess the role of AhRR in immunometabolism and mitochondrial function, WT and AhRR-deficient FLiM cells were analyzed in a Seahorse Mitochondrial Stress test. The cellular metabolic state of macrophages is linked to their activation state. Alternatively activated macrophages are often characterized by an enhanced oxidative metabolism, whereas classically activated macrophages rely on glycolysis [57]. Here, we could show that AhRR-deficient FLiM cells had higher basal and maximal respiration rates than WT FLiM cells, implying that lack of AhRR expression shifts macrophages towards alternative activation (Fig. 2D, S2B). Enhanced oxidative metabolism can lead to an increase in reactive oxygen species (ROS) generation as a side product of the electron transport chain. Recent data implied that intestinal IELs of AhRR^-/-^ mice showed increased intracellular ROS [36]. In accordance with these results, AhRR^-/-^ FLiM cells also had elevated ROS levels after stimulation with Zymosan (Fig. 2E). In line, analysis of the mitochondrial membrane potential using Tetramethylrhodamine (TMRM) staining showed a higher accumulation of the dye in mitochondria of AhRR^-/-^ cells, implying enhanced mitochondrial activity in AhRR-deficient FLiM cells (Fig. 2F, S2C). Additionally, we analyzed bone marrow-derived macrophages (BMMΦ) and primary liver macrophages from WT and AhRR^-/-^ mice. In line with the results shown for FLiM cells, AhRR-deficient BMMΦ and primary macrophages isolated from the liver exhibited an enhanced maximal and basal respiration rate (Fig. 2G-H, S2D-E). Altogether, these data show that AhRR-deficient macrophages have an altered immunometabolic phenotype with increased mitochondrial capacity, leading to enhanced oxidative phosphorylation and ROS production.

### 2.3 Mice lacking AhRR solely in myeloid cells do not phenocopy global AhRR-deficient mice but show reduced diet-induced hepatosteatosis

Following these results, we hypothesized that myeloid cells, known to influence the metabolism of hepatocytes [43–45] and adipocytes [46–48] via cellular cross-talk, could be responsible for the reduced weight gain and reduced liver steatosis observed in global AhRR-deficient mice. To directly address this question, we generated AhRR^fl/fl^ mice (Fig. S3A) and crossed them with LysM^Cre^ mice [58] to generate mice deficient for AhRR in myeloid cells. Deletion efficiency in macrophages was proven via qPCR (Fig. S3B). Different from our original hypothesis, after challenging AhRR^fl/fl^LysM^Cre^ mice with HFD for 14 weeks, we did not observe differences in either body weight or fat mass compared to Cre-negative controls, although moderately lower glucose levels could be detected in blood (Fig. 3A-C). Moreover, and in contrast to AhRR^-/-^ mice, AhRR^fl/fl^LysM^Cre^ mice had decreased oxygen consumption at 23°C (Fig. 3D). In line with these results, no differences in weight, morphology or adipogenic and pro-fibrotic markers were observed in the WATg (Fig. S4A-D). Nevertheless, mRNA levels of the gene encoding fatty acid synthase (*Fasn*), linked to visceral adipose tissue accumulation, along with mRNA levels of monocyte chemoattractant protein-1 (*Mcp1*) were significantly decreased in AhRR^fl/fl^LysM^Cre^ mice compared to AhRR^fl/fl^ control mice (Fig. S4C, E). In addition, in the HFD-fed groups, the liver weight was significantly lower (-20.86%) in AhRR^fl/fl^LysM^Cre^ mice compared to control HFD mice (Fig. 3E). Histological analysis revealed reduced steatosis and lower expression levels of the fatty acid binding protein 4 (*Fabp4*), commonly associated with metabolic-associated fatty liver disease (MAFLD) [59] (Fig. 3F-G). Similarly, the more moderate occurrence of fibrosis assessed by Sirius Red staining and reduced expression of the pro-fibrotic marker *Col1a1* (Fig. 3H-I) indicated a reduced fatty liver disease in AhRR^fl/fl^LysM^Cre^ mice compared to AhRR^fl/fl^ HFD-fed mice. To further corroborate these findings, we analyzed the levels of plasma ALT after HFD feeding. ALT levels were increased in AhRR^fl/fl^ control mice, whereas they remained unchanged in AhRR^fl/fl^LysM^Cre^ mice upon HFD feeding (Fig. 3J). Altogether, these data demonstrate that the lack of AhRR in myeloid cells protects from severe diet-induced hepatosteatosis and slows down progression of liver disease.

**Figure 3:**
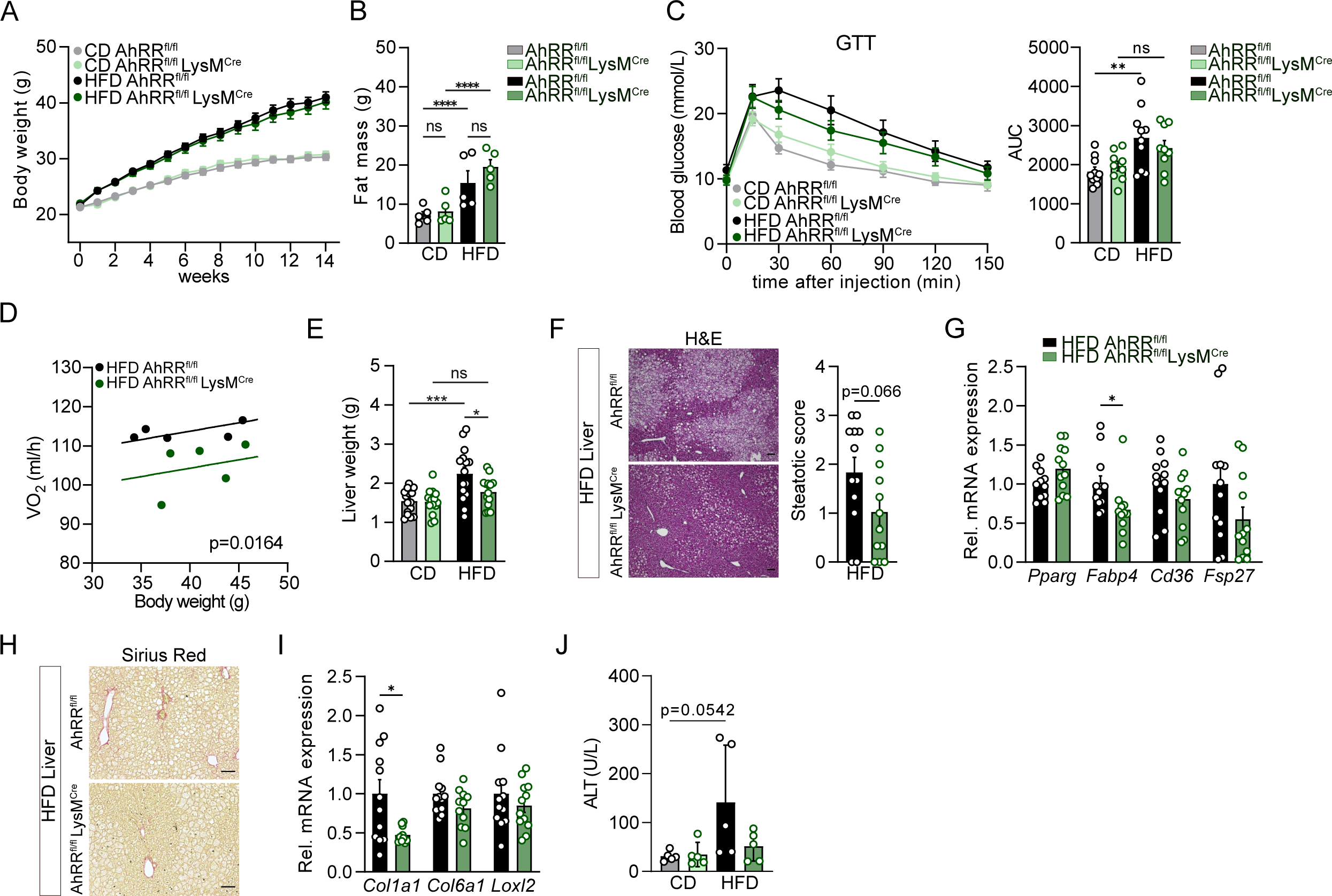
Lack of AhRR in myeloid cells protects from diet-induced liver steatosis. (**A-J**) WT (AhRR^fl/fl^) mice and mice lacking AhRR in myeloid cells (AhRR^fl/fl^LysM^Cre^) were fed either a CD or a HFD over the course of 14 weeks. (**A**) Body weight; n=14-16 (CD) and n=15 (HFD). (**B**) Total body fat mass assessed by NMR; n=5. (**C**) Blood glucose over the course of 150 min (left) and AUC quantification (right); n=9-10. (**D**) ANCOVA of oxygen consumption rate of HFD mice; n=5. (**E**) Liver weight, n=14-16. (**F**) Representative H&E (left panel) and semiquantitative analysis of the steatotic score (right panel) of HFD liver samples. Scale bars, 100 µm; n=12. (**G**) Relative mRNA expression levels of lipid-related markers in HFD liver samples, n=12. (**H**) Representative Sirius Red staining of HFD liver samples. Scale bars, 100 µm. (**I**) Relative mRNA expression of pro-fibrotic markers in HFD liver samples; n=12. (**J**) ALT concentration in plasma of AhRR^fl/fl^ and AhRR^fl/fl^LysM^Cre^ mice fed either CD or HFD for 14 weeks; n=5. *p < 0.05, **p ≤0.01, ***p ≤ 0.001. Significance was determined using unpaired two-tailed t-tests (F, G, and I) and one-way ANOVA with Tukey’s multiple-comparison test (B, C, E and J). Data are mean ± s.e.m.

### 2.4 Lack of AhR in myeloid cells aggravates diet-induced hepatosteatosis and fibrosis

Next, we analyzed if the protective effects seen in AhRR^fl/fl^LysM^Cre^ mice could be recapitulated in AhR^fl/fl^LysM^Cre^ mice, as global deficiency of both, AhRR and AhR conferred protection from DIO and MAFLD. Remarkably, like AhRR^fl/fl^LysM^Cre^ mice, AhR^fl/fl^LysM^Cre^ mice were not protected from diet-induced weight gain following 14 weeks of HFD feeding (Fig. 4A). In line with these results, no differences in WATg weight, expression of adipogenic markers (*Fabp4, Pparg, Adipoq, Cd36, and Fasn*), and serum cholesterol levels were observed (Fig. 4B-D). Strikingly, histological analysis of WATg revealed a high accumulation of F4/80-positive cells in AhR^fl/fl^ LysM^Cre^ mice compared to AhR^fl/fl^ mice (Fig. 4E). Similarly, AhR^fl/fl-^LysM^Cre^ mice showed increased collagen deposition, accompanied with a significantly higher fibrosis score and increased *Col1a1* and *Col6a1* mRNA levels (Fig. 4F-G). In line with these results, histological analysis showed increased hepatosteatosis, whereas the liver weight was not changed (Fig. 4H-I). Although *Pparg* levels were significantly decreased, its direct target *Fsp27*, which prevents lipid mobilization and promotes intracellular lipid storage [60], was increased by 81.5% on average (Fig. 4J). Moreover, the liver of AhR^fl/fl^LysM^Cre^ mice appeared more fibrotic and depicted increased *Col1a1* and *Col6a1* mRNA expression levels (Fig. 4K). In line, we observed an average increase of 37.6% in serum ALT levels (Fig. 4L). Altogether these data indicate that loss of AhR in myeloid cells severely affects the consequences of DIO accelerating the progression of MAFLD and aggravating visceral adipose tissue fibrosis.

**Figure 4:**
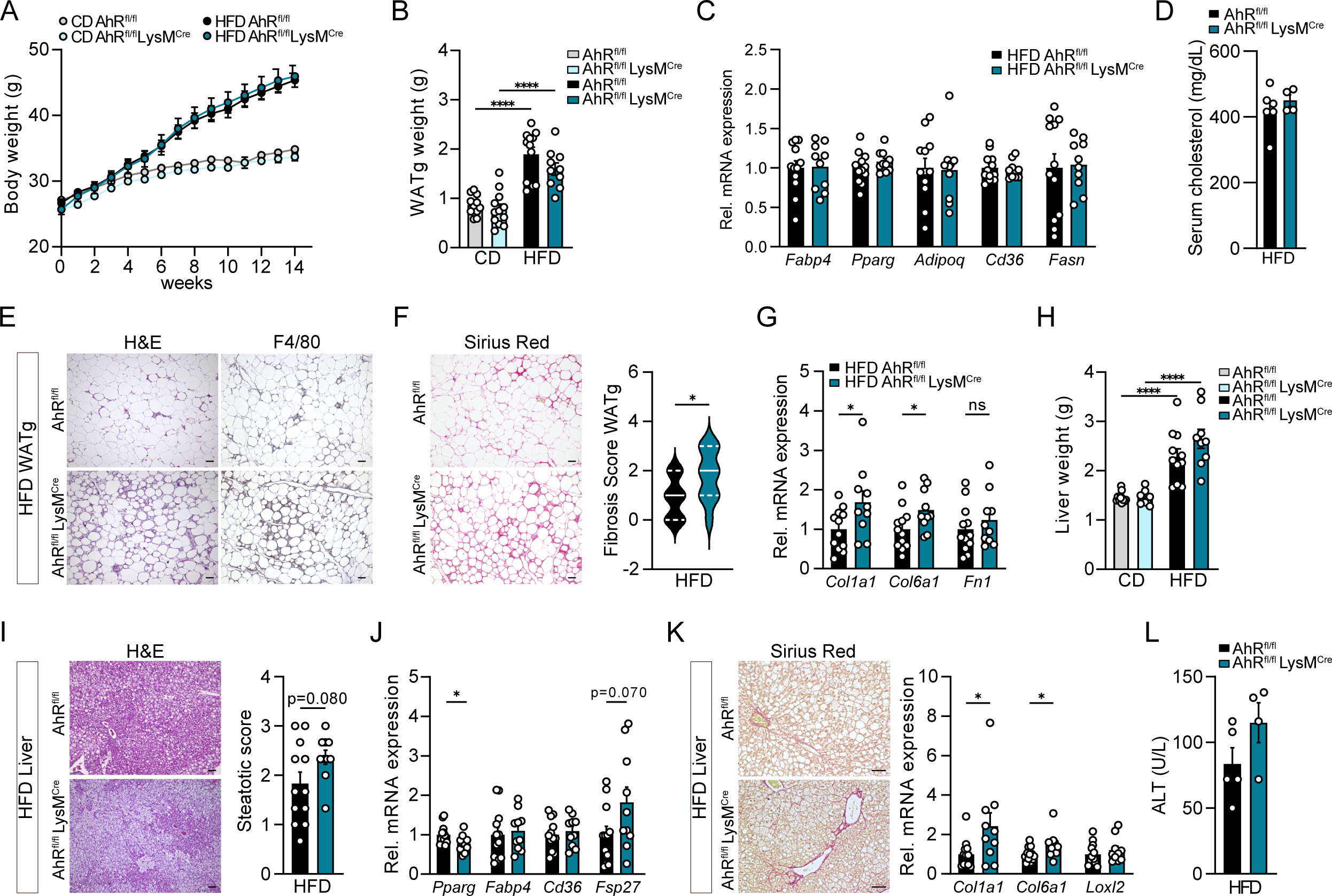
Lack of AhR in myeloid cells aggravates hepatic steatosis and induces fibrosis in liver and adipose tissue. (**A-L**) WT (AhR^fl/fl^) mice and mice lacking AhR in myeloid cells (AhR^fl/fl^LysM^Cre^) were fed either a CD or a HFD over the course of 14 weeks. (**A**) Body weight; n= 10-12. (**B**) WATg weight in grams; n= 10-12. (**C**) Relative mRNA levels of adipogenic markers in WATg of HFD mice; n= 10-12. (**D**) Total serum cholesterol; n=4-6. (**E**) Representative H&E (left panel) and F4/80 (right panel) staining of the WATg of HFD mice. Scale bars, 100 µm. (**F**) Representative Sirius Red staining (left panel) and semiquantitative analysis of the fibrosis score (right panel) of the WATg of HFD mice. Scale bars, 100 µm; n=10-11. (**G**) Relative mRNA levels of pro-fibrotic markers in WATg of HFD mice; n=10-12. (**H**) Liver weight; n=10-12. (**I**) Representative H&E stainings (left panel) and semiquantitative analysis of the steatotic score (right panel) of HFD liver samples. Scale bars, 100 µm; n=10-12 (**J**) Relative mRNA expression levels of lipid-related markers in the liver of HFD mice; n=10-12. (**K**) Representative Sirius Red staining of liver sections. Scale bars, 100 µm (left panel). Right, relative mRNA expression levels of pro-fibrotic markers in the liver of HFD mice; n=10-12. (**L**) Serum ALT concentration of AhR^fl/fl^ and AhR^fl/fl^LysM^Cre^ mice fed a HFD for 14 weeks; n=4-5. *p < 0.05, **p ≤0.01, ***p ≤ 0.001. Significance was determined using unpaired two-tailed t-tests (C, D, F, G, I, J, K, and L) and one-way ANOVA with Tukey’s multiple-comparison test (B and H). Data are mean ± s.e.m.

## 3. Discussion

In the last decades, obesity has become one of the leading risk factors associated with poor cardiometabolic health, diabetes, dyslipidemia, and cancer [61,62]. It is known that the AhR links recognition of environmental stimuli to the homeostasis of metabolic organs [63,64] as well as overall metabolic health and development of obesity [19,50]. In addition, AhR deficiency can protect mice from DIO associated with ameliorated hepatic steatosis and increased energy expenditure [8,19]. Here, we analyzed the influence of the AhRR on the development of DIO and liver damage. We previously described that AhRR expression is mainly detected in hematopoietic cells of barrier organs and their draining lymph nodes [32] but the role of the AhRR in macrophage programming and its influence on metabolic diseases was still unclear. Unexpectedly, global deficiency of the AhRR protected mice from DIO, similar to the effects observed in AhR^-/-^ mice. This was accompanied by an increase in whole-body energy expenditure and diminished hepatosteatosis, thereby identifying AhRR as a potential regulator of metabolic health. Alongside improved glucose tolerance and insulin sensitivity, AhRR^-/-^ mice showed decreased body weight and fat mass. As we observed a reduced size of adipocytes in WATg of global AhRR^-/-^ mice, it cannot be excluded that the AhRR to some extent also controls adipocyte metabolism directly. More likely, however, a cross-talk of several cell types, in particular immune cells and non-hematopoietic parenchymal or stromal cells accounts for the ameliorated DIO in global AhRR^-/-^ mice.

A possible explanation for the finding that AhR- and AhRR-deficient mice show a similar phenotype regarding the development of DIO is the differential regulation and a stronger cell type-specific expression of the AhRR compared to AhR. A similar scenario has been observed in the context of dextran sodium sulfate (DSS)-induced intestinal inflammation where AhR- as well as AhRR deficiency aggravated susceptibility to inflammatory bowel disease. In the intestine, the AhRR is expressed in various immune cell subtypes but not in epithelial cells, whereas AhR expression is of particular importance in intestinal epithelial cells controlling the expression of CYP enzymes [32,65,66]. Thus, we hypothesize that AhRR deficiency ameliorates DIO because of enhanced AhR activity in certain cell types [32], an effect that may also be achieved by higher ligand availability. In line, Choi et al. showed that dietary supplementation of the AhR ligand precursor I3C regulated adipogenic and thermogenic gene programs such that DIO was reduced [51]. We confirmed this effect of decreased weight gain and ameliorated hepatosteatosis in WT mice fed I3C-supplemented HFD but observed no further reduction in weight gain and steatosis by I3C in AhRR-deficient mice. Thus, AhRR deficiency has a stronger effect than I3C supplementation.

Macrophages, which are key regulators of tissue homeostasis and repair, express the AhRR [32]. Previous studies showed that macrophage metabolism influences their phenotype and plasticity to adapt to different environments in health and disease, thus enabling specific effector functions [67]. Therefore, we assessed how AhRR deficiency affects cellular programming in macrophages. Here, we observed increased oxidative phosphorylation in conjunction with higher mitochondrial membrane potential in AhRR-deficient macrophages, indicating an overall enhanced mitochondrial activity. Interestingly, the AhR signaling pathway has been identified as a regulator of mitochondrial homeostasis in the liver [68]. ROS are generated as a side product of oxidative phosphorylation in the mitochondrial electron transport chain [69] or as a result of CYP enzymatic activity [70], which is enhanced in the absence of the AhRR. In accordance with the study by Panda et al., reporting an increased ROS production in AhRR-deficient IELs [36], we detected elevated macrophage ROS production and alterations in the level of plasma TCA cycle intermediates after HFD feeding. In line with alterations in mitochondrial metabolism, transcriptomic analysis of FLiM cells revealed an enrichment of Myc and mTORC1 pathways known to regulate mitochondrial biogenesis and function besides other cellular processes [71,72]. Altogether, our data underline the importance of AhRR in regulating mitochondrial activity.

Further, genes associated with *de novo* lipogenesis, such as *Acly* encoding ATP citrate lyase and *Acaca* encoding Acetyl-CoA carboxylase, both essential for fatty acid synthesis, were enriched in AhRR-deficient FLiM cells. It has been shown that changes in *Acaca* affect the bioenergetic profile of macrophages and influence their response to pro-inflammatory stimuli [73]. Moreover, *Acly* expression is associated with increased ROS and prostaglandin E2 production and has been suggested to increase phagocytosis in TLR-activated BMMΦ [74]. Wculek et al. revealed that macrophages with high demand for lipid handling rely on mitochondrial respiration to meet their metabolic need [75]. Thus, the OXPHOS capacity of macrophages directly correlates with their functional capacity to handle high lipid burden. Consequently, we hypothesize that in the absence of AhRR expression, the overall enhanced expression of *de novo* lipogenesis genes and increased mitochondrial activity contribute to alternative activation and elevated macrophage ROS levels.

Despite the observed alterations in metabolic programming of AhRR-deficient macrophages, analysis of mice with a conditional loss of AhRR in myeloid cells only partially recapitulated the phenotype of global AhRR-deficient mice in the DIO model. Thus, AhRR^fl/fl^LysM^Cre^ mice were not protected from developing DIO and metabolic syndrome but showed protection from severe hepatosteatosis and obesity-associated liver injury. In contrast, AhR^fl/fl^LysM^Cre^ mice revealed an exacerbation of liver disease following 14 weeks of HFD. It was shown before that Kupffer cells, liver-resident macrophages, interact with hepatocytes and thereby shape the tissue environment, regulating hepatic lipid metabolism and the liver’s response to HFD [43,76,77]. Moreover, macrophages in the liver are key players in MAFLD development and are responsible for the efferocytosis of lipid-laden hepatocytes [45,78]. Thus, through deletion of the AhRR in myeloid cells, the progression of MAFLD was slowed down, although the mice still exhibited signs of metabolic syndrome. It should be noted that besides macrophages, neutrophils are also affected by LysM^Cre^-mediated deletion [58]. As granulocytes do not express *Ahr* nor *Ahrr* [79], it is unlikely, however, that neutrophils account for the observed phenotype. Taken together, our results suggest AhRR as a rheostat of macrophage metabolism and function conferring a more protective and lipid-tolerant status in the liver.

Liver fibrosis often occurs as a consequence of chronic liver damage and is positively correlated with higher mortality rates [80]. Interestingly, we observed a significant decrease in the expression levels of the pro-fibrotic marker *Col1a1* in the liver of HFD-fed AhRR^fl/fl^LysM^Cre^ mice. In contrast, AhR^fl/fl^LysM^Cre^ mice displayed higher expression of *Col1a1* and *Col6a1*, alongside exacerbated Sirius Red staining. The role of AhR in hepatic fibrosis, however, is controversially discussed. While global AhR-deficient mice spontaneously developed periportal liver fibrosis [81], TCDD-induced activation of the AhR has also been described to induce hepatic fibrosis [82,83]. In contrast, the provision of another AhR ligand was shown to ameliorate CCl_4_-induced fibrosis. It has been reported that global AhR deficiency promotes the activation of hepatic stellate cells (HSC), key players in the development of hepatic fibrosis [84]. Liver macrophages are the main regulators of HSC function, producing pro-fibrogenic cytokines, which support survival of activated HSC [85]. Activated HSC secrete chemokines attracting macrophages, and consequently aggravate the progression of liver fibrosis. Of note, loss of AhR expression in HSC but not in hepatocytes appeared to be responsible for the development of periportal fibrosis [84]. Similarly, macrophages are also involved in adipose tissue fibrosis, stimulating extracellular matrix (ECM) accumulation [86], but also clearing excessive ECM through collagen uptake and degradation [87]. Here we demonstrate that loss of AhR in macrophages worsens the development of HFD-induced fibrosis, while AhRR^fl/fl^LysM^Cre^ mice show reduced collagen deposition. Therefore, the expression of AhR in macrophages and the regulation of AhR signaling via AhRR both play a role in the development of diet-induced fibrosis depending on external stimuli.

Besides the differential role of AhR and AhRR in macrophages in the context of DIO and liver damage, the functional importance of the AhR strongly differs between cell types and stage of development. For example, early developmental ablation of the AhR either globally or in endothelial cells causes failure of ductus venosus closure in the liver [88] which may affect diet-induced liver damage. As a result, considering its restricted cell type-specific expression [32,36], the AhRR may represent a more promising target for potential pharmacological therapy of metabolic diseases than the AhR itself.

## 4. Materials and methods

### 4.1 Animals and Experimental animal procedures

All mice were bred and maintained in individually ventilated cages under specific pathogen–free conditions. AhRR^-/-^, AhRR^fl/fl^, AhRR^fl/fl^LysM^Cre^, and AhR^-/-^ mice [89] were bred at the animal facility of the LIMES, Bonn, Germany. AhR^fl/fl^LysM^Cre^ mice were bred at the animal facility of the IUF, Düsseldorf, Germany. In all experiments with myeloid cell-specific conditional knockout mice, the LysM^Cre^-knockin allele was heterozygous (Cre/+). WT and littermate mice served as controls. All mice were on a C57BL/6JRcc background.

Seven to nine-week-old male mice were used for the experiments and were bred according to German guidelines for animal care. For *in vivo* experiments, mice were age-matched within 2 weeks and randomly assigned to treatment groups. All experiments were performed according to German and Institutional guidelines for animal experimentation (permits: AZ 84-02.04.2015.A043, 81-02.04.2019.A262 and 81-02.04.2017.A435) and were approved by the government of North Rhine-Westphalia (Germany).

For dietary intervention studies, mice were exposed to a purified diet or with 2 g/kg I3C-supplemented control diet (CD/ CD+I3C, 13 kJ% Fat, Cat No.: D12450 LS ssniff EF) or a purified or with 2 g/kg I3C-supplemented High fat diet (HFD/ HFD+I3C, 60 kJ% fat, Cat. No.: D12492 ssniff EF) ad libitum. The body weight was assessed weekly. After 14 weeks of feeding, the mice were analyzed.

### 4.2 Generation of AhRR^fl/fl^ mice

To generate AhRR-floxed mice, loxP recognition sequences were introduced in the second and third intron of the *Ahrr* gene, flanking the third exon, which contains the bHLH domain of the AhRR essential for DNA binding [90,91]. An frt-flanked neomycin resistance gene was inserted behind the 5’ loxP site to allow for positive selection of the targeted ES cells. After electroporation of the targeting vector into HM-1 ES-cells (Sv129/OlaHsd), selected clones were screened by PCR and Southern blot using a 3’flanking probe. Successfully recombinant clones were injected into C57BL/6JRcc blastocysts. Germline transmission was proven by PCR analysis. Mice were backcrossed to C57BL/6JRcc for 8 generations.

The frt-flanked neomycin resistance cassette was deleted by directly introducing flippase (flp) mRNA into fertilized oocytes of homozygous AhRR-floxed (AhRR^fl/fl^) mice. StemMACS Flp Recombinase mRNA (130-106-769, Miltenyi Biotec GmbH) was introduced by electroporation (7 square wave pulses of 30 V, 3ms, 0,1s pause) using a GenePulser Xcell electroporator (Biorad) and a 1 mm cuvette (Biorad). Differentiated 2-cell stage embryos were transferred into pseudopregnant females BALB/cAnNCrl (50%) C57BL/6NCrl (50%). Offspring was screened by PCR and sequencing of the locus for successful removal of the neomycin cassette. The targeting strategy is depicted in Fig. S3A.

To generate AhRR-deficient mice, AhRR^fl/fl^ mice were bred to Cre-deleter mice. Deletion of the third exon of the *Ahrr* was demonstrated by PCR analysis and the Cre allele was removed by backcrossing the mice to C57Bl/6JRcc mice.

To delete AhRR in myeloid cells, AhRR^fl/fl^ mice were crossed to LysM^Cre^ mice [58]. The deletion efficiency was evaluated by qPCR. A control fragment, giving rise to a signal in both AhRR^fl/fl^ and LysM^Cre^-deleted strains (primers House 350 BP) and a fragment spanning the deleted allele after Cre recombination (primers dCRE) was amplified from genomic DNA of BMMΦ of AhRR^fl/fl^ and AhRR^fl/fl^LysM^Cre^ mice. A standard reaction using WT and AhRR-deficient genomic DNA was included.

Primers:

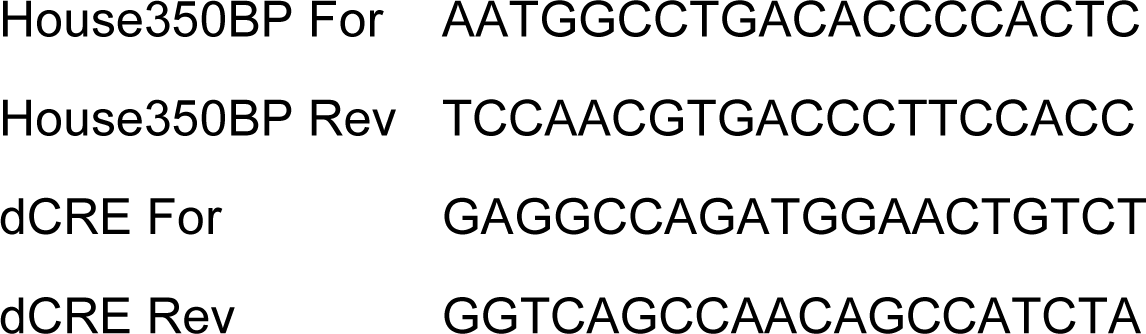

### 4.3 Glucose and Insulin Tolerance Test

Glucose tolerance tests (GTT) or insulin tolerance tests (ITT) were performed at week 12 of feeding. A drop of blood from the nicked tail vein was used to determine the basal glucose concentration using AccuChec Instant test strips (C216796165-IMP, Roche) after six hours of fasting for GTT and without fasting for ITT. Then, mice were injected intraperitoneally with 0.25 mg/L glucose at a dose of 2g/kg of body weight or 0.09375 U/mL insulin at a dose of 0.75 U/kg of body weight, respectively. Blood glucose levels were subsequently measured at 15, 30, 60, 90, 120, and 150 min post-injection.

### 4.4 Measurement of Serum and Plasma Parameters

Piccolo Express Chemistry Analyzer (Abaxis Inc.) and Reflotron Plus Analyzer (Roche Diagnostics) were used to measure the concentration of blood serum and plasma parameters. Blood samples were collected into EDTA tubes from the *vena facialis* of experimental animals. Serum was tested for Alanine Aminotransferase (ALT/GPT, Ref 10745138), and Cholesterol (Ref 10745065) levels using respective test strips (Roche Diagnostics) following manufacturer’s instructions. Using the Piccolo analyzer, the Lipid Panel Plus (Ref 400-1030) was obtained.

### 4.5 Macrophage Staining

Paraffin-embedded samples from WATg were sectioned and deparaffinized. Antigen retrieval was performed with 10 mM Sodium Citrate Buffer (pH 6) at 85°C for 50 min. After treatment with BloxAll Blocking Solution (VectorLabs, Cat.-No. SP-6000), samples were blocked with 2.5% goat serum and stained with rat anti-F4/80 (clone A3-1, Santa Cruz, Cat.-No. sc-59171) and subsequently HRP-coupled goat anti rat IgG secondary antibody (ImmPRESS Goat Anti-Rat IgG Polymer Kit Peroxidase, VectorLabs, Cat.-No. MP7404). The HRP substrate reaction was performed with DAB (ImmPACT DAB Substrate Kit, Peroxidase, Cat.-No. SK-4105), samples were counterstained with Hemalaun, and mounted with VectaMount Permanent Mounting Medium. Images were acquired with a Keyence BZ-9000 Microscope (Keyence Corporation).

### 4.6 Oil Red O Staining

Livers were fixed in 4% paraformaldehyde (PFA) for at least four hours, dehydrated by treatment with ascending concentrations of D-Sucrose for at least two hours per solution, and embedded in Tissue Tek. 10 µm sections were washed in Formalin, rinsed in 60% Isopropanol, and stained in freshly prepared Oil Red O staining solution (5 g/L in Isopropanol). The samples were counterstained with Hemalaun and mounted with Kaiser’s glycerol gelatin with Phenol. Randomized images were obtained using a Keyence BZ-9000 microscope.

Quantitative analysis was performed using Fiji (Rasband, W.S., ImageJ, U. S. National Institutes of Health, https://imagej.nih.gov/ij/, 1997-2018). Thereby, the tissue was distinguished from background signal in the pictures and the mean intensity of red signal in the tissue was calculated. Subsequently, the Optical Density (OD) was calculated from the values of mean intensity, normalizing the mean intensity on the maximal intensity (value of maximal intensity=255) with the following formula: OD=log(Maximal Intensity/Mean Intensity). The script for quantification is provided upon request.

### 4.7 Culture of Bone Marrow-derived Macrophages (BMMΦ)

For isolation of BMMΦ, the femur and tibiae of mice were flushed with sterile PBS. The bone marrow cells were cultured and differentiated at 37°C and 5% CO_2_ in RPMI medium supplemented with 10 – 15% supernatant of L929 cells, 10% FCS, penicillin/ streptomycin, 50 µM β-mercaptoethanol and 2 mM L-Glutamine. Fresh medium was added on day 4 of the culture. After seven days, the cells were harvested and used for downstream readouts.

### 4.8 Isolation of Liver Macrophages

Liver macrophage isolation was performed as previously described [92]. In brief, livers were perfused with EDTA and Heparin in HBSS and subsequently with 50 mg/mL collagenase and CaCl_2_ in Williams E medium via the portal vein. After perfusion, the liver was removed from the diaphragm and put into a beaker glass to shake the organ. The resulting cell suspension was transferred into a 50 mL reaction tube and centrifuged for 2 min at 20 g. The supernatant of the first centrifugation, containing the Kupffer cells, was re-centrifuged at 500 g for 10 min. The supernatant was subjected to Magnetic Activated Cell Sorting (MACS) using CD11b magnetic microbeads (Order No. 130-093-634, Miltenyi Biotec) according to the manufacturer’s instructions and over LS columns after red blood cell lysis. After washing, the eluted cells were plated to a Seahorse 96 well cell plate in a range from 100,000 to 200,000 cells/well.

### 4.9 Generation of Fetal Liver Macrophage (FLiM) cell lines

FLiM cells were generated as described in Fejer et al. 2013 [54]. Briefly, single cell suspensions of fetal livers of E13.5 mouse embryos were prepared and cultured in T75 flasks in FLiM medium (RPMI 1640, 10 % FCS, 2 mM L-Glutamine, 100 U/mL Penicillin/Streptomycin, 50 µM β-mercaptoethanol with 2% supernatant of granulocyte-macrophage colony-stimulating factor (GM-CSF) transfected X63Ag8–653-cells). When a stable proliferation rate was reached, cells were split 1:20 based on confluence every 6-7 days, while fresh medium was added 2-3 days after splitting.

### 4.10 Seahorse Assay (Mito Stress Test)

One day before the assay (BMMΦ) or on the day of the assay (liver macrophages), 1-2 x 10^5^ cells were seeded in XF96 cell culture microplates (Seahorse Bioscience). The Sensor Cartridge was prepared according to the manufacturer’s instructions. One hour prior to the assay, growth medium was discarded and replaced by Seahorse Assay XF RPMI assay medium (pH 7.4), and the plate was incubated for 60 min at 37 °C. Mito Stress Test inhibitor stock solutions were adjusted to appropriate concentrations and pipetted into the ports of the Sensor Cartridge (Port A: 20 µL of 10 µM Oligomycin; Port B: 22 µL of 40 µM FCCP; Port C: 25 µL of 5 µM Rotenone/ Antimycin A). A Seahorse Bioscience XFe96 Analyzer was used to measure extracellular acidification rates (ECAR) and oxygen consumption rates (OCR) every 3 min. After Seahorse measurement, cell numbers were normalized using the CyQuant NF Cell Quantification kit (ThermoFisher Scientific).

### 4.11 ROS Assay

FLiM cells were seeded in FLiM medium at a concentration of 1-3 x 10^5^ cells/mL in a white 96 well plate at 37°C 5% CO_2_. After 24 h the plate was centrifuged at 400 g for 5 min, the medium was discarded and 170 µL/well fresh FLiM medium was added. Directly before measuring, 30 µL/well of the Luminol mix containing 80 U/mL Horseradish peroxidase (Merck) and 8 mM Luminol (AppliChem Life Science) in Borate buffer (0.2 M H_3_BO_3_ 0.02 M Na_2_B_4_O_7_x10H_2_O) was added including either 0.4 mg/mL Zymosan (Sigma Aldrich) or PBS for control samples. ROS production was measured in relative luminescence units (RLU) via a luminometer (Berthold Technologies) for a period of 30-60 min. Area under the curve was calculated.

### 4.12 Flow cytometric analysis of mitochondrial membrane potential using Tetramethylrhodamine Methyl Ester (TMRM)

0.75 x 10^6^ FLiM cells were seeded in FLiM medium in a six well plate and incubated overnight at 37°C 5% CO_2_. The Mitoprobe TMRM kit (ThemoFisher) was used according to manufacturer’s instructions. Prior to flow cytometric analysis, cells were harvested, washed with PBS, and then resuspended in 500 µL PBS before measuring TMRM intensity using the PE bandpass filter of a FACSCanto (BD Biosciences).

### 4.13 Metabolic cages and determination of body composition

#### Energy expenditure

Oxygen consumption was measured for 120 s per cage during 24 h using Phenomaster (TSE Systems). All mice were maintained on a daily cycle of 12-hours light (06:00–18:00 hours) and 12-hours darkness (18:00–06:00 hours), at 23 ± 1°C and were allowed free access to chow and water.

#### Body composition analysis

Body composition was analyzed using a table Bruker Minispec.

### 4.14 H&E and Sirius Red staining

WATg and liver were fixed in PBS containing 4 % PFA overnight and dehydrated using ethanol. Dehydrated tissues were embedded in paraffin. Paraffin-embedded samples from WATg and liver were cut in 5 µm sections. Sections were deparaffinized and hydrated to distilled water.

#### Haematoxylin and eosin (H&E)

Sections were stained with Hemalaun for 3 min and rapidly washed with distilled water. Staining with eosin was performed right after by incubating the tissue sections in eosin for 3 min and washing with distilled water. Following dehydration, samples were mounted in Entellan (107960, Sigma-Aldrich) and dried overnight. Images were acquired with a Keyence BZ-9000 Microscope (Keyence Corporation). Blinded manual scoring of liver steatosis was performed using the grading from Sethunath et al. [93].

#### Sirius Red

Sections were covered and incubated for 60 min with Picrosirius Red solution (SRS500, ScyTek Laboratories). Slices were quickly rinsed in a 0.5 % acetic acid solution (Cat #AAD, ScyTek), followed by a quick rinse in 100 % ethanol. Samples were embedded in Euparal (Art. No. 7356.1, Roth) and dried overnight. Images were acquired with a Keyence BZ-9000 Microscope (Keyence Corporation). The Fibrosis Score of stained WATg samples was assessed according to Lassen et al. [94].

### 4.15 RNA isolation and qRT-PCR

RNA from WATg and liver was isolated as described [95]. Final concentration of RNA was quantified using a Nanodrop Spectrophotometer. First-strand cDNA was synthesized from 1000 ng of total RNA using mixture of oligo(dT)12–18 primers and Superscript reverse transcriptase (Thermofisher).

mRNA expression was assessed by qRT-PCR using a Real-Time system CFX96 (BioRad) and Absolute qPCR SYBR Green ROX mix (ThermoFisher). Quantification of mRNA levels was performed based on the crossing point values of the amplification curves using the second derivative maximum method. mRNA expression levels were normalized to *Hprt* and were displayed as fold change relative to samples of control mice used as calibrator.

The primer sequences are shown in the following table:

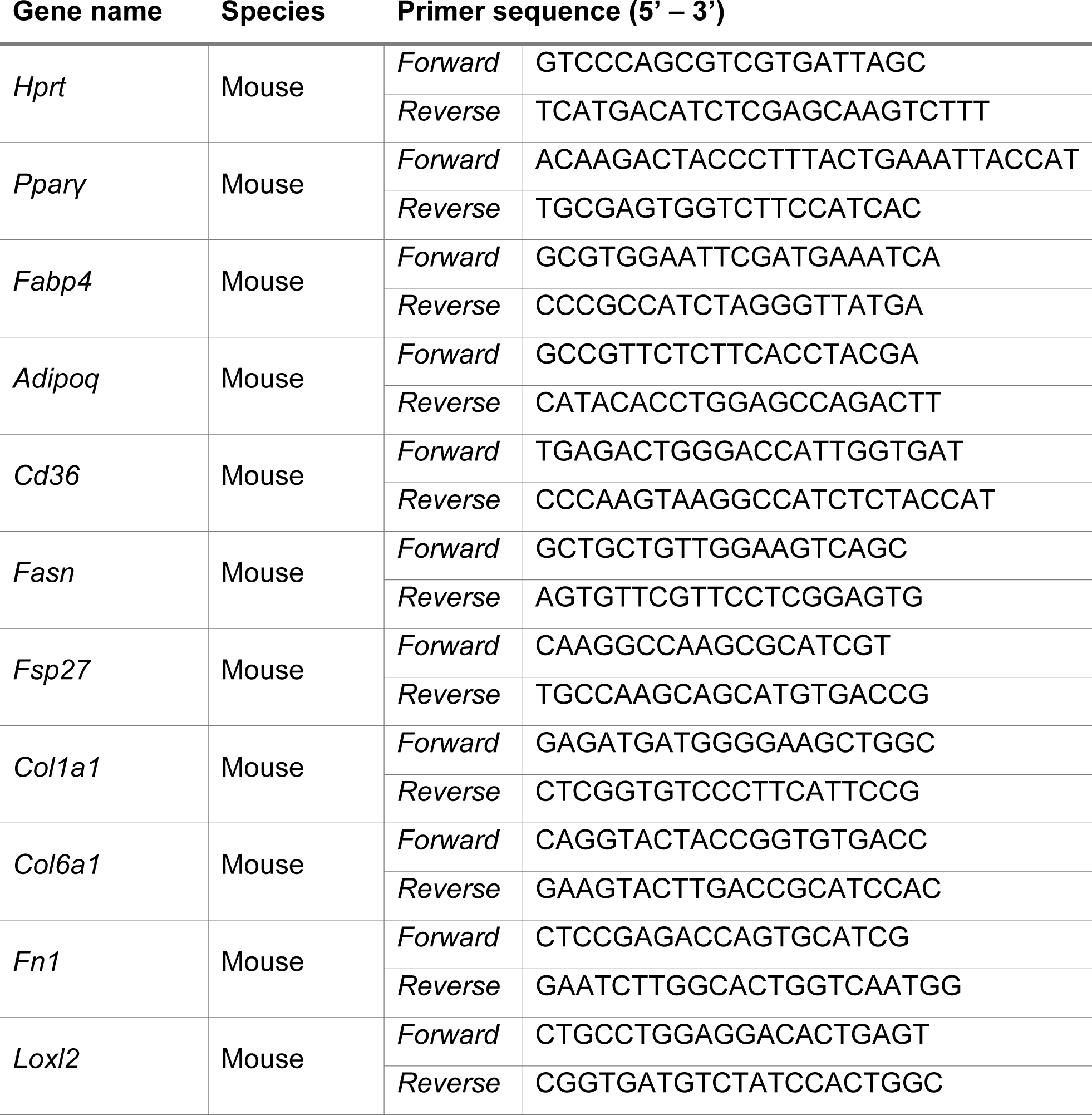
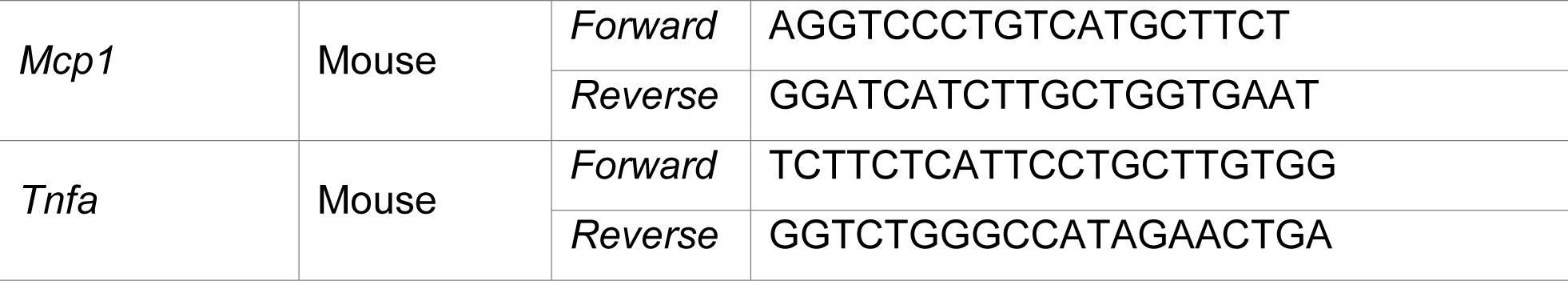

### 4.16 Metabolite extraction from plasma samples

Extraction was performed as previously described [96,97]. Briefly, 20 µL plasma sample was added to 180 µL ice-cold methanol:H_2_O (8:1) containing 2 µg/mL D6-pentanedioic acid (C/D/N Isotopes, Quebec, Canada) as internal standard. The mixture was vortexed at 1400 rpm for 10 min at 4 °C and centrifuged at 17,000 g for 10 min at 4 °C. 80 µL supernatant was transferred into glass vials compatible with gas chromatography and then dried under vacuum at 4 °C in a CentriVap Concentration System (Labconco, Kansas City, Missouri).

### 4.17 GC-MS measurement and data analysis

GC-MS measurement of relative metabolite levels was performed as described [98], using an Agilent 7890B gas chromatograph equipped with a 30 m DB-35ms and 5 m Duraguard capillary column (Agilent, Santa Clara, California) for separation of derivatized metabolites, and an Agilent 5977B MSD system (Agilent) for measurement of metabolites. Briefly, dried metabolite extracts were derivatized with equal amounts of methoxylamine (20 mg/mL in pyridine, both Sigma-Aldrich, Steinheim, Germany) and MSTFA or MTBSTFA (Restek Corporation, Bellefonte, Pennsylvania) before injection into the GC-MS system. Measurements were carried out in either full scan or selected ion mode. Processing of chromatograms and relative quantification of metabolites were performed using the Metabolite Detector software [99].

### 4.18 Transcriptome analysis

Cells were lysed using Qiazol (Qiagen) for RNA isolation, and RNA was extracted using the miRNeasy kit (Qiagen), following the manufacturer’s instructions. Polyadenylated RNA was then enriched from total RNA using Oligo-dT-coupled magnetic beads. The RNA was fragmented and transformed into double-stranded cDNA libraries using the TruSeq RNA Sample Preparation Kit v2 (Illumina), following manufacturer’s protocol. cDNA fragments, favoring 200 bp, were selected and purified.

Sequencing was done on a HiSeq1500 device in a 75bp single-end run, after cluster generation on a cBot station (Illumina). Raw sequencing reads underwent quality control with fastQC 0.11.9 (https://www.bioinformatics.babraham.ac.uk/projects/fastqc/) and were summarized using MultiQC v1.14 (https://multiqc.info/). The 75 bp single-end reads were trimmed using fastp v0.20.0 and aligned to the murine mm10 vM32 reference genome from GENCODE using STAR v2.7.10b (https://github.com/alexdobin/STAR, [100]) with -- quantMode GeneCounts. Aligned reads sorted by coordinates were indexed using samtools v1.16.1. The alignment process was managed using SnakeMake 7.20.0 (https://snakemake.github.io/, [101]). Subsequent data analysis was conducted in R version 4.2.2 RStudio 2023 09.0 Build 303, mainly utilizing the DESeq2 version 138.3 R package [102]. Raw counts were imported into R, and a DESeq object was created using the DESeqDataSetFromMatrix function. Genes with fewer than 10 total counts were omitted, leaving 16,111 genes for analysis. Size factors and dispersions were estimated per gene using the default settings for DESeq2. Unwanted variations, like technical variance from donors, were addressed in the normalization process or removed using surrogate variable analysis (sva) [103]. The rlog-transformed expression values, adjusted for surrogate variables identified by sva, were processed with the “removeBatchEffect” function from the limma package [104]. Differential expression analysis was conducted using DESeq2, incorporating SV1 and SV2 into the design model for comparing AhRR^-/-^ samples to WT controls. This analysis, which applied a fold change (FC) threshold of 1.5, utilized the ‘apeglm’ method for shrinkage of estimates and the Independent Hypothesis Weighting (IHW) approach for multiple testing correction, identified 312 genes as upregulated and 360 genes as downregulated at an adjusted p-value cutoff of 0.05. Gene set enrichment analysis (GSEA) [105] was carried out on the transcriptome data for all Hallmark gene sets [106], comparing AhRR^-/-^ versus WT controls. Only gene sets with an adjusted p-value under 0.05 (Benjamini-Hochberg correction) were considered and visualized. Differentially expressed genes were filtered for transcription factors [107], surface and secretome markers from the Human Protein Atlas [108], and metabolism genes based on literature. The top 20 genes from each category were visualized in a heat map. Sequencing data associated with this paper has been submitted to NCBI’s GEO repository under the accession number GSE254369.

## Acknowledgements

We are grateful to Babette Martiensen for technical help. This work was supported by the Deutsche Forschungsgemeinschaft (DFG, German Research Foundation) through SFB 1454 (Project No. 432325352) (to I.F., K.H., M.D.B., A.P. and C.T.), EXC 1023 (Project No. 194445620) (to I.F.), EXC 2151 (Project No. 390873048) (to I.F.), and TRR 333/1 – 450149205 (to I.F. and A.P.); the University of Bonn through an Argelander grant to F.G., as well as the Jürgen Manchot Foundation (to I.F. and H.W.).

## Author Contributions

Conceptualization, F.J.G., F.G., A.P., C.E., H.W., I.F. and L.R.-S.; methodology, F.J.G., F.G., Y.M., B.N., I.I., K.W., Z.A., W.H., K.H., T.U., H.W., L.R.-S.; software, T.U.; validation, F.J.G., K.H., T.U., C.E., H.W., L.R.-S.; formal analysis, F.J.G., T.U. and L.R.-S.; investigation F.J.G., F.G., Y.M., B.N., K.K., M.G., M.M., I.I., P.H., S.K.L., K.W., W.H., K.Hä.; resources, C.T., Z.A., K.H., M.D.B., A.P., C.E., H.W. and I.F.; data curation, F.J.G., T.U. and L.R.-S.; writing—original draft preparation, F.J.G, Y.M., H.W., I.F. and L.R.-S.; writing—review and editing, H.W., I.F. and L.R.-S.; visualization, F.J.G., Y.M., H.W. and L.R.-S.; supervision, A.P., C.E., H.W., I.F. and L.R.-S; funding acquisition, F.G., K.H., A.P., C.E., H.W. and I.F.

## Figure Legends

**Figure S1:**
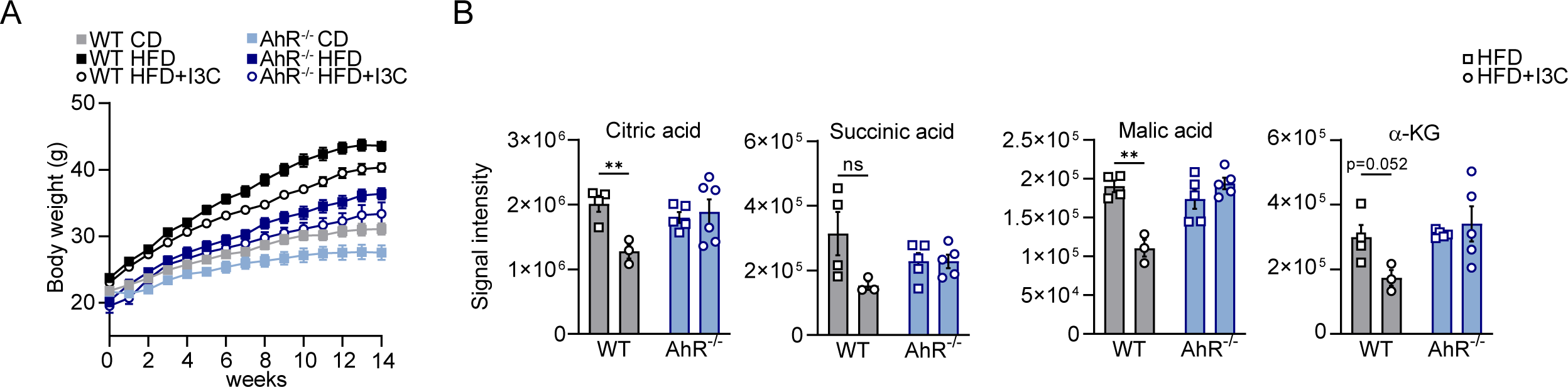
AhR-deficient mice are resistant to diet-induced obesity. (**A**) Body weight of wild-type (WT) and AhR-deficient mice fed a CD or HFD ± I3C for 14 weeks. (**B**) Citric acid, succinic acid, malic acid and a-KG levels in the plasma of WT and AhR^-/-^ HFD ± I3C mice; n=3-5. *p < 0.05, **p ≤0.01, ***p ≤ 0.001. Significance was determined using multiple unpaired two-tailed t-tests. Data are mean ± s.e.m.

**Figure S2:**
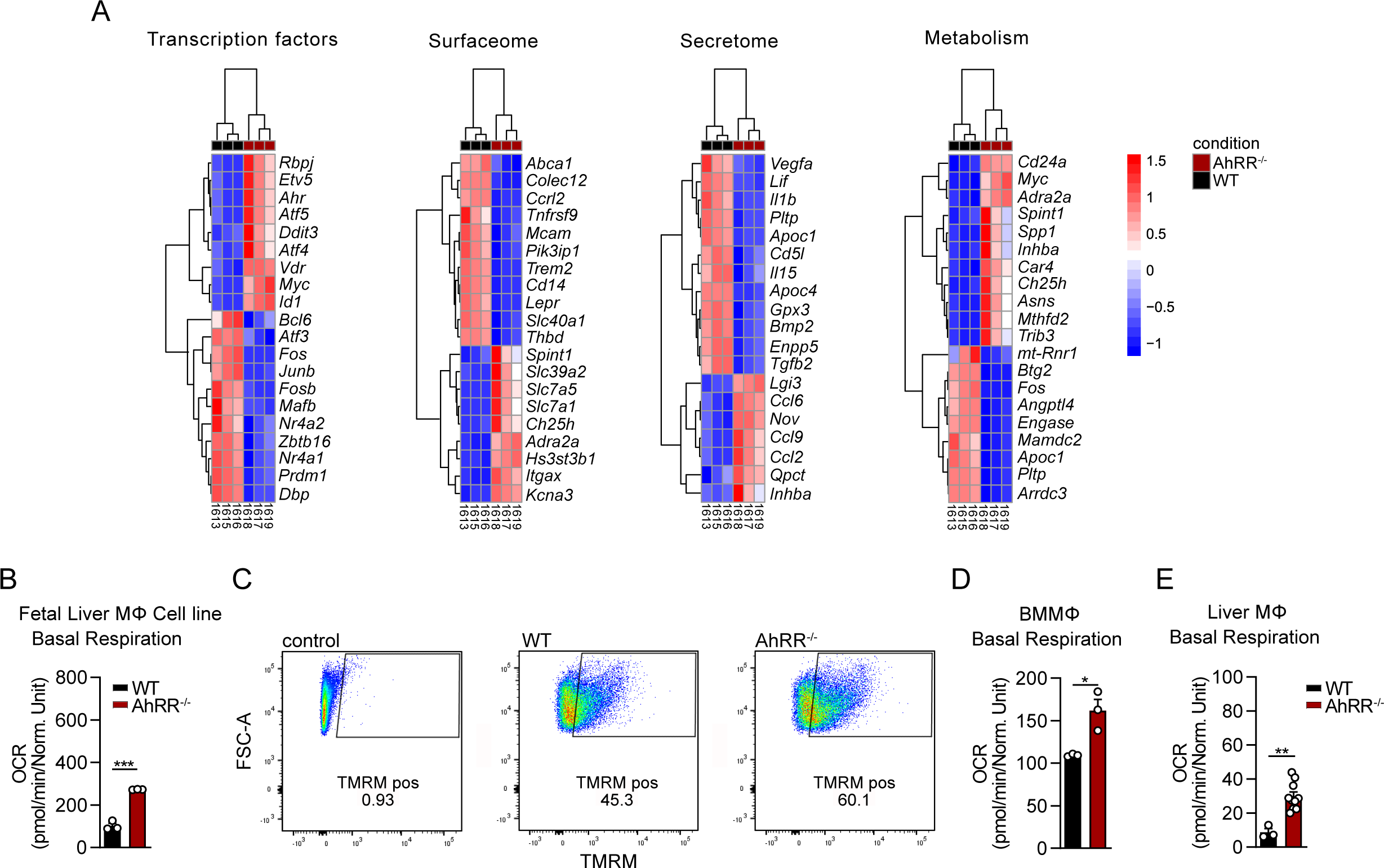
AhRR-deficient macrophages display an altered metabolic and gene expression profile. (**A**) Differentially expressed genes filtered by known transcription factors, surfaceome, secretome, and metabolism, n=3. (**B**) Basal respiration rate after mitochondrial stress test analysis in fetal liver macrophage (FLiM) cell lines of WT and AhRR-deficient mice; n=3. (**C**) Representative flow cytometry data of assessment of the mitochondrial membrane potential in WT and AhRR-deficient FLiM cells. (**D-E**) Basal respiration rate after mitochondrial stress test analysis in (**D**) bone marrow derived macrophages (BMMΦ) and (**E**) primary liver macrophages isolated from AhRR^-/-^ and WT mice. For D, n=3 and for E, n=3 (WT), n=10 (AhRR^-/-^). Seahorse data show mean ± s.e.m. of three technical replicates from one representative experiment. *p < 0.05, **p ≤0.01, ***p ≤ 0.001. Significance was determined using unpaired two-tailed t-tests. Data are mean ± s.e.m.

**Figure S3:**
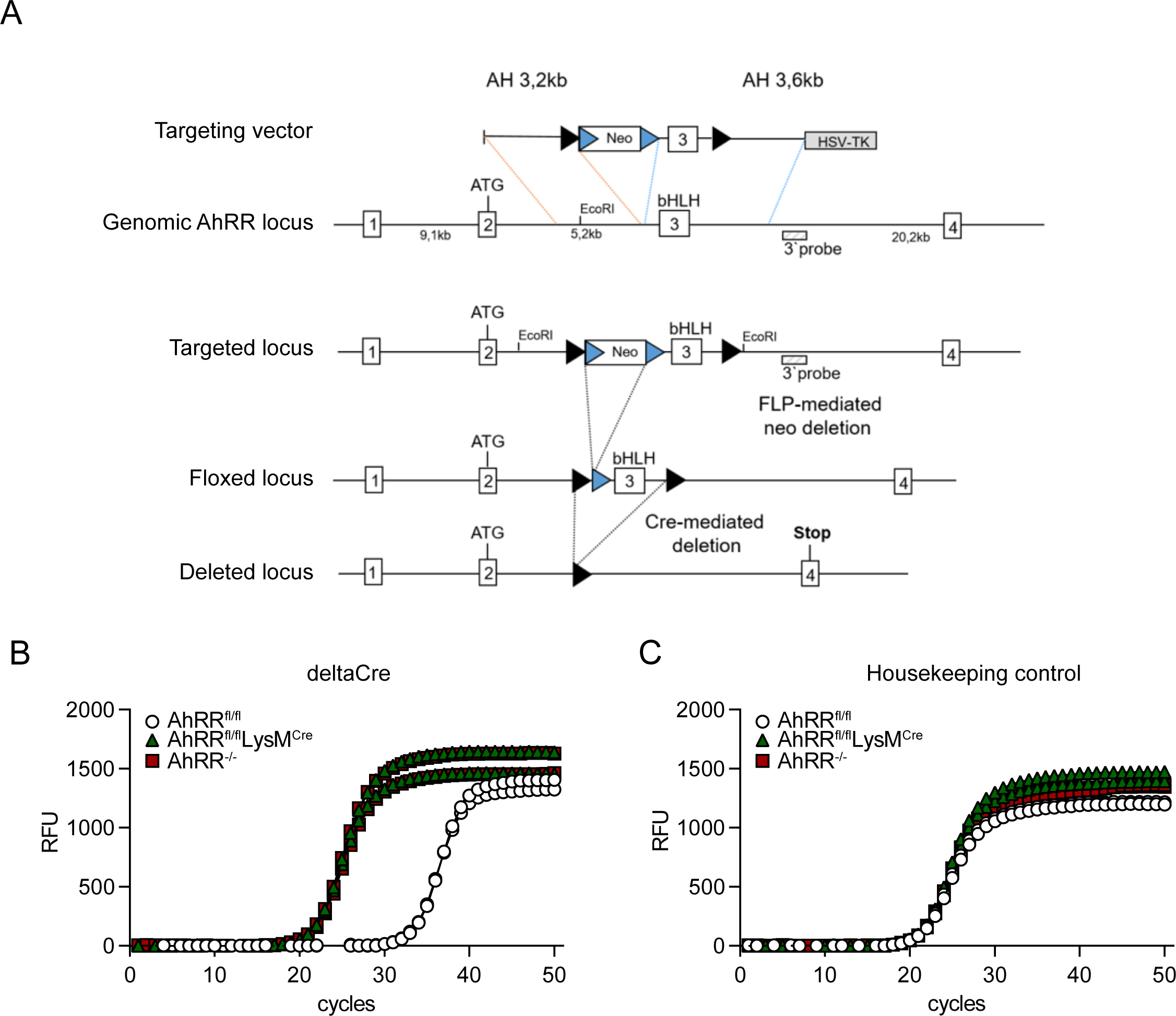
Generation of AhRR-floxed mice. (**A**) Targeting strategy for generation of AhRR-floxed mice. LoxP recognition sequences were introduced in the second and third intron of the *Ahrr* gene, flanking the third exon. A frt-flanked neomycin cassette was placed behind the 5’loxP site and was removed from AhRR^fl/fl^ mice by delivering flippase (flp) mRNA into fertilized oocytes. (**B**) To generate mice with a deletion of AhRR in myeloid cells, AhRR^fl/fl^ mice were crossed to LysM^Cre^ mice. Deletion efficiency was estimated by qPCR analysis of AhRR^-/-^, AhRR^fl/fl^LysM^Cre^ and AhRR^fl/fl^ BMMΦ showing the amplification product of the deleted *Ahrr* locus (deltaCre) occuring simultaneously in AhRR^-/-^ and AhRR^fl/fl^LysM^Cre^ BMMΦ but significantly later in AhRR^fl/fl^ control BMMΦ. A control amplification product of a house keeping gene appeared in all three genotypes (right). Shown are two independent replicates for each genotype.

**Figure S4:**
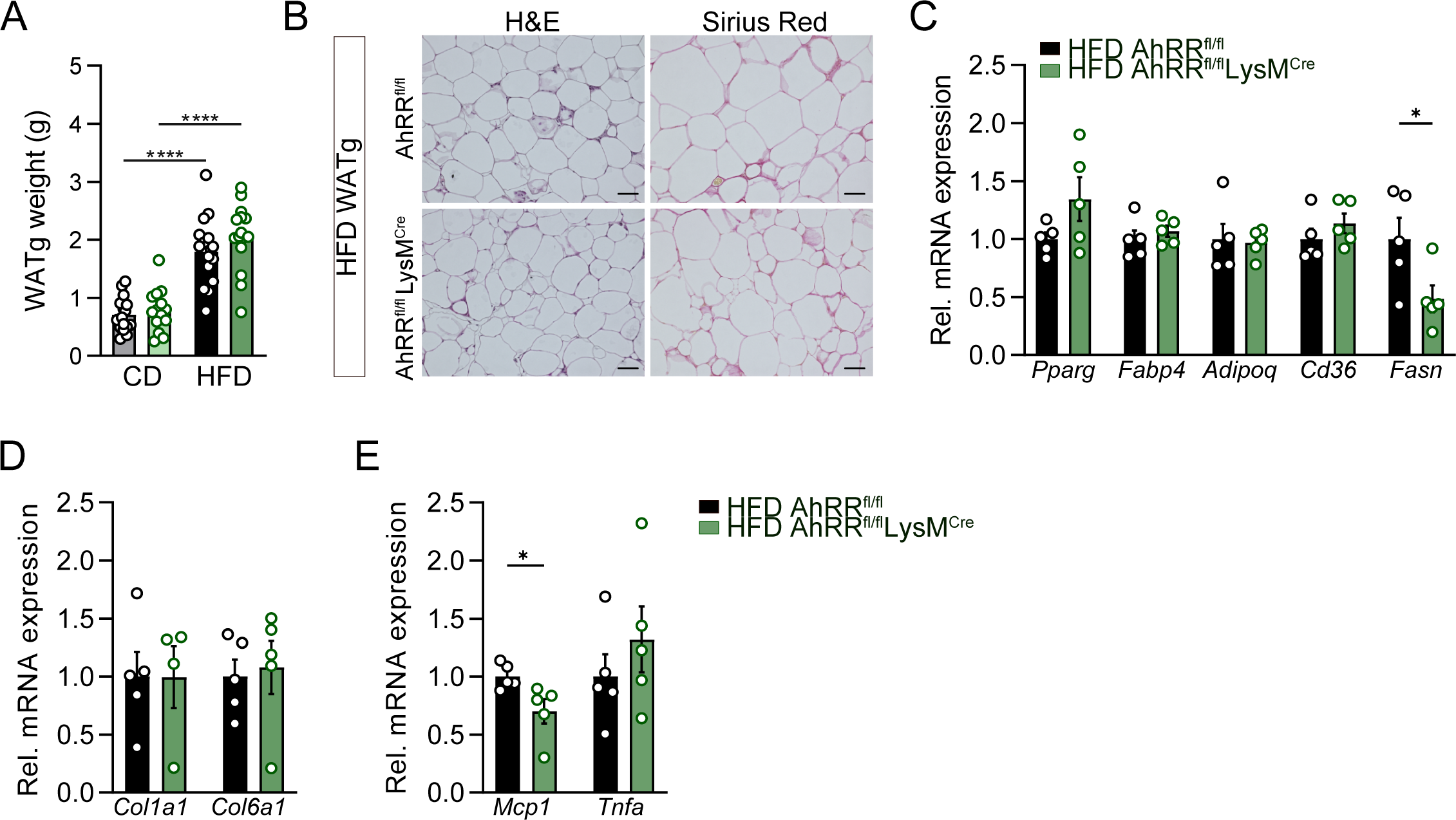
Analysis of WATg of WT and AhRR^fl/fl^LysM^Cre^ mice after HFD. (**A-F**) WT (AhRR^fl/fl^) and AhRR^fl/fl^LysM^Cre^ mice after 14 weeks of CD or HFD feeding. (**A**) WATg weight; n=14-16. (**B**) Representative H&E (left panel) and Sirius Red (right panel) staining of WATg sections of WT and AhRR^fl/fl^LysM^Cre^ mice after HFD. Scale bars, 100 µm. (**C-E**) Relative mRNA expression of (**C**) adipogenic markers, (**D**), pro-fibrotic markers, and (**E**) pro-inflammatory markers in the WATg of WT and AhRR^fl/fl^LysM^Cre^ mice after 14 weeks feeding HFD, n=5. *p <0.05, **p ≤0.01, ***p ≤ 0.001. Significance was determined using unpaired two-tailed t-tests (C, D, and E) and one-way ANOVA with Tukey’s multiple-comparison test (A). Data are mean ± s.e.m.

